# Beyond the Matrix: Rethinking Antibiotic Tolerance in CF Biofilms Using 3D Models

**DOI:** 10.1101/2025.07.19.665652

**Authors:** Goodness Ogechi Osondu-Chuka, Stephan Schandl, Andrea Scheberl, Olivier Guillaume, Aleksandr Ovsianikov, Erik Reimhult

## Abstract

Chronic lung infections in cystic fibrosis (CF) patients are associated with *Pseudomonas aeruginosa* biofilms exhibiting high antibiotic tolerance with no clear explanation. We investigate the role of the biofilm matrix in this antibiotic tolerance using 3D biofilm models based on acetylated alginate and DNA, mimicking mucoid biofilms. Printed from these bioinks seeded with *P. aeruginosa* (PAO1), these models support robust microcolony formation as observed in vivo and enable high-throughput assessment of antibiotic diffusion and efficacy. Surprisingly, antibiotic diffusion is not significantly impeded by acetylation or DNA incorporation. Despite this, bacterial tolerance increases tremendously upon encapsulation in alginate. Acetylation further enhances tolerance, particularly to tobramycin, ciprofloxacin, and colistin. The addition of DNA mitigates this effect in a drug-specific manner. While mucoid biofilms, in contrast to the biofilm models, show significant retardation of antibiotic penetration, they also get saturated with all tested antibiotics within 20 h. This demonstrates that direct interaction with alginate or DNA does not explain the slow diffusion of antibiotics in mucoid *P. aeruginosa* biofilms. Our findings challenge the view that diffusion limitation or antibiotics binding by biofilm exopolysaccharides dominate biofilm resilience and highlight the need to target matrix-induced bacterial adaptation in the development of antibiofilm therapies.

## 1. Introduction

Cystic fibrosis is a genetic disorder that predominantly affects the lungs, causing the production of thick, viscous mucus that impairs normal respiratory function and fosters an environment conducive to chronic bacterial infections. ^[1, 2]^ In CF patients, *P. aeruginosa* (PA) emerges as a predominant pathogen responsible for persistent pulmonary infections. ^[3–5]^ This microorganism forms biofilms, which serve as a defensive mechanism against both antimicrobial treatments and the host’s immune system. ^[6, 7]^ The presence of biofilms, characterized by their dense extracellular matrix (ECM), facilitates the persistence of infections that become impossible to eradicate. ^[8]^ The chronic nature of PA infections in CF patients is further exacerbated by the emergence of mucoid bacteria variants, which overproduce alginate, a key exopolysaccharide (EPS) that enhances biofilm stability and antibiotic tolerance. ^[9, 10]^

The EPS plays a vital role in PA’s ability to withstand antimicrobial treatment, with alginate, Pel, and Psl acting as protective and structural components. ^[11–13]^ Notably, PA secrete acetylated alginate, a modification that has been suggested to contribute to antibiotic tolerance by altering the biofilm’s physicochemical properties compared to pristine alginate. ^[14–16]^ In addition, extracellular DNA (eDNA), which originates from bacterial and host cell lysis or active secretion, contributes to biofilm integrity and interacts with exopolysaccharides such as Psl to enhance structural strength and antimicrobial tolerance. ^[17–20]^ The strong polyanions alginate and eDNA are thought to play an instrumental role in inhibiting the diffusion of predominantly cationic antibiotics, preventing them from efficiently reaching bacteria within the biofilm,^[21–25]^ but the weakly anionic Pel and cationic Psl were also implicated in biofilm PA antibiotic tolerance.^[26–28]^ While alginate and eDNA have long been regarded as physical barriers that hinder antibiotic diffusion,^[25, 29–32]^ the extent to which biofilm tolerance arises directly from these physicochemical impediments and their direct contributions to adaptive bacterial responses (e.g., transcriptional and metabolic changes) remains poorly resolved. Additionally, although the individual contributions of alginate, Pel, Psl, and eDNA to biofilm architecture are increasingly well-characterized, their dynamic interactions with antibiotics within a relevant 3D context remain poorly understood. This highlights the need for innovative, matrix-defined biofilm models to more accurately explore the mechanisms underlying PA’s drug tolerance in biofilms.

To address the limitations of traditional biofilm research, the development of 3D biofilm models has emerged as a significant advancement. ^[33, 34]^ Unlike conventional *in vitro* systems, such as microtiter plate-based and Calgary biofilm models, 3D biofilm models more accurately replicate the structural and functional complexities of *in vivo* biofilms, including the spatial organization, nutrient gradients, and diffusion barriers that influence bacterial behavior and antibiotic responses. ^[34, 35]^ These models could enable the study of mucoid biofilms in a controlled environment, providing critical insights into the mechanisms of biofilm resilience and potential therapeutic targets. To bridge the gap between traditional biofilm models and the complex *in vivo* environment of the CF lung, we developed a 3D biofilm model that incorporates key biofilm components, including synthetically acetylated alginate and DNA.^[16, 36]^ Alginate acetylation is characteristic of PA mucoid biofilms, but its impact on biofilm properties is poorly investigated.^[16]^ We demonstrated by varying the degree of acetylation that achieving the 36% degree of acetylation in native biofilm alginate was critical for mimicking the physicochemical properties of mucoid biofilm matrices using alginate hydrogel models. ^[16, 36]^Still, its influence on antibiotic tolerance is unknown. Our novel model provides a more physiologically relevant platform for studying the mechanisms of antibiotic tolerance in mucoid PA biofilms, while ensuring knowledge of its detailed composition and physicochemical properties.

We use this model to investigate the role of alginate acetylation and DNA in PA biofilm-associated antibiotic tolerance using 3D biofilm models for a battery of antibiotics clinically used against PA—tobramycin (TOB), gentamicin (GEN), ciprofloxacin (CIP), colistin (COL), meropenem (MER), and aztreonam (AZT). By examining how alginate acetylation and DNA influence drug diffusion and bacterial persistence, we seek to enhance our understanding of biofilm-associated antibiotic tolerance and ultimately contribute to the development of more effective therapeutic strategies for chronic CF infections. In particular, we address whether antibiotic tolerance in biofilms is a direct consequence of antibiotic interaction with these dominant matrix components or if it is an emerging property triggered by encapsulation.

## 2. Results

### 2.1 Mimicking Bacterial Alginate for a More Accurate CF Infection Model

To replicate the *in vitro* biofilm microenvironment of CF-associated infections, it was essential to synthetically modify readily available alginate produced by brown seaweed algae with controlled degrees of acetylation within the reported range of mucoid PA. ^[37–39]^ With a method of selective acetylation, modified alginate with acetylation predominantly on its mannuronic acid (M) units was developed.^[16]^ Alginate in PA biofilms has been characterized to have a degree of acetylation of their M-units of 22-40%.^[40]^ To fabricate our biofilm models, alginate with a degree of acetylation of 36 % (**Figure S1a**) was used as it closely mimics acetylation of alginate in mucoid PA biofilms. While the native mucoid alginate extracted from PA exhibited a higher molecular weight (∼3×) than the seaweed-derived alginate used in this study, mucoid PA alginate samples considerably vary between strains and even within a strain under different environmental conditions.^[16, 41, 42]^ Such variability can compromise reproducibility across experiments. In contrast, selectively acetylated seaweed-derived alginate enables precise control over acetylation and block composition, allowing consistent modeling of alginate-mediated antibiotic tolerance. Based on our previously published results, the synthesized acetylated alginate closely matched the physicochemical and mechanical properties of mucoid PA alginate before and after tobramycin treatment. ^[16, 36]^ We also demonstrated that these modified alginates closely mimic the rheological, diffusion, and antibiotic-binding properties of native mucoid alginate. ^[16, 36]^

### 2.2 PA Growth and Distribution in Biofilm Models

Biofilm models were molded as ∼7 mm disks or printed as ∼600 µm beads from alginate with bacteria suspensions crosslinked by Ca^2+^, washed, and transferred to TSB. Incubating the biofilm models at 37 °C resulted in the formation of bacterial microcolonies within 24 h, as seen in **Figure 1**.

**Figure 1.**
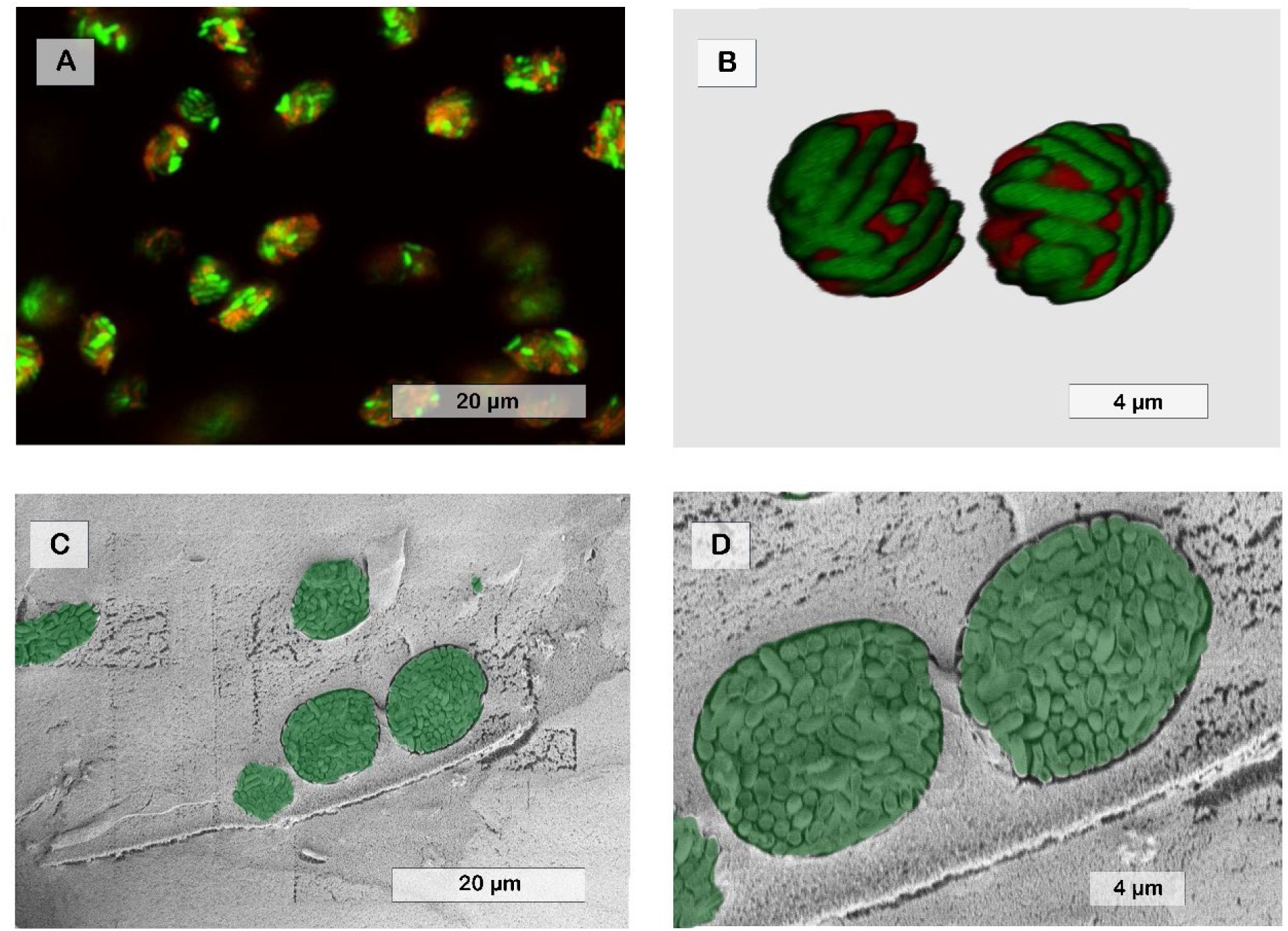
Shape and spatial distribution of PAO1 microcolonies in acetylated alginate biofilm models showing round morphologies expanding from individual seeded cells. A) CLSM image of PAO1 microcolonies distributed in acetylated alginate biofilm model matrix stained with SYTO-9 and PI. B) CryoSEM micrograph depicting cross-sections of acetylated alginate biofilm models, highlighting sectioned microcolonies (manually coloured green). The biofilm models were incubated for 24 h at 37 °C.

The size and the spatial distribution of these aggregates within the alginate beads were analyzed using a Confocal Laser Scanning Microscope (CLSM) and a Cryogenic Scanning Electron Microscope (CryoSEM). The PAO1 microcolonies were homogenously distributed throughout the matrix. They ranged in size from approximately 7 µm to 15 µm and contained ∼185 ± 1.3 – 476 ± 8.1 cells per colony, respectively. This distribution of microcolonies demonstrated the successful original dispersion of PA in the printed biofilm model, with each dispersed cell expanding into a separate microcolony. Viability assays confirmed that the recovered cells remained viable after 24 h (**Figure 1** and **Figure S2 – S3**), with the population increasing by approximately four orders of magnitude, from an initial count of ∼25 × 10^9^ CFU ml ^−1^ to ∼15 × 10^13^ CFU ml ^−1^.

### 2.3 Impact of Alginate Acetylation on Antibiotic Diffusion in Biofilm Model

The ability of antibiotics to diffuse through biofilms is a crucial factor influencing their efficacy. We first investigated the impact of alginate acetylation on antibiotic diffusion by measuring the diameter of the zone of inhibition (ZOI) for *E. coli* growth produced by antibiotics diffusing through each biofilm model in the format described in **Figure S4**.^[43]^

The diffusion of most of the antibiotics through non-acetylated and acetylated biofilm models was at most weakly affected compared to an agarose model used as a reference (**Figures 2** and **S4**). The diffusion rates of aminoglycosides exhibited slight variations but were, however, not statistically significant (p ≥ 0.05) (**Figure S4**).

**Figure 2:**
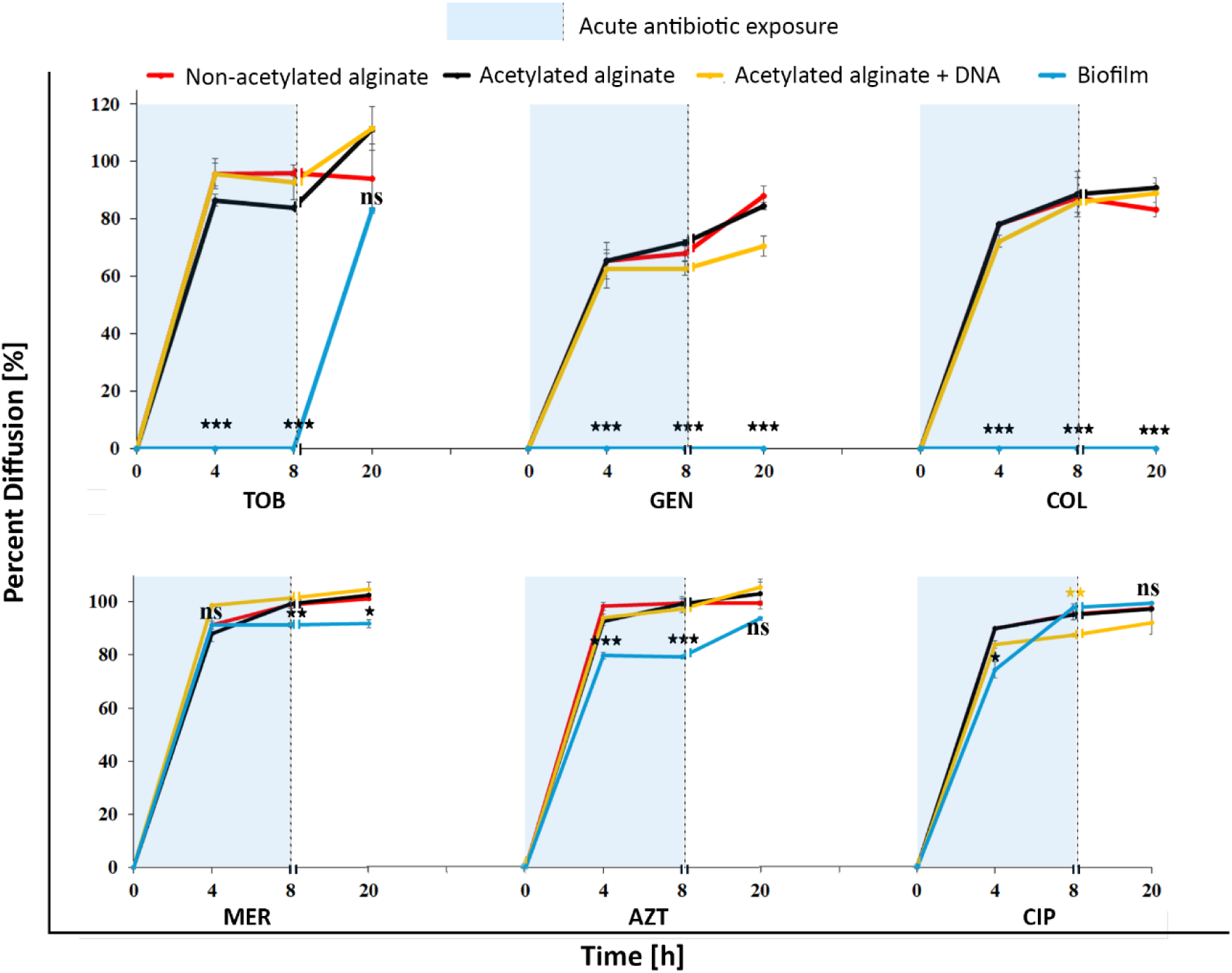
Line graphs representing diffusion percentages of selected antibiotics through the disk *in vitro* biofilm models and mucoid PA biofilms at different time points, normalized to diffusion through agarose disks, and quantified by a zone of inhibition assay.^[43]^ All models showed similar antibiotic diffusion, while biofilms (blue line) significantly hindered antibiotic diffusion of tobramycin, gentamicin, and colistin. Statistical significance between mucoid PA biofilm and other *in vitro* biofilm models [* p ≤ 0.05, ** p ≤ 0.01, *** p ≤ 0.001, ns = non-significant, **\*\*** p ≤ 0.01 (only for acetylated alginate + DNA)]. Statistical significance for all *in vitro* biofilm model comparisons is provided in **Supplementary Figure S4**.

Colistin, ciprofloxacin, aztreonam, and meropenem demonstrated progressive increases in diffusion from 4 to 20 h in both models, with no statistical significance (p ≥ 0.05). In summary, over 20 h, all antibiotics except for gentamicin and colistin reached equilibrium levels in both acetylated and non-acetylated alginate biofilm models.

### 2.4 Effect of DNA on Antibiotic Diffusion in Acetylated Alginate Models

To assess the effect of extracellular DNA (eDNA) on antibiotic tolerance and binding to cationic antibiotics, DNA (from Calf thymus) was mixed with acetylated alginate and cast as biofilm models. The antibiotic diffusion through this model was compared with diffusion through an agar model containing DNA. **Figures 2** and **S4** demonstrated that adding DNA did not significantly affect tobramycin diffusion (p ≥ 0.05) at any time point, and the tobramycin concentrations reached equilibrium levels by 20 h in both models. (Statistical significance for all *in vitro* biofilm model comparisons is provided in **Supplementary Figure S4**.) In contrast, for gentamicin, the presence of DNA significantly reduced diffusion at 8 and 20 h (p ≤ 0.05). For ciprofloxacin, the presence of DNA significantly reduced diffusion only at 8 h (p ≤ 0.05), with no significant difference at earlier or later time points. Similarly, colistin’s diffusion was significantly reduced at 4 h (p ≤ 0.05) but not at 8 h or 20 h. In the case of meropenem, the addition of DNA led to a significant increase (p ≤ 0.05) in diffusion at 4 h. There was no significant difference at later timepoints. Finally, aztreonam reached equilibrium diffusion levels after 20 h in all models with no statistical difference between the tested groups.

In summary, we show that the impact of DNA on antibiotic diffusion is antibiotic-specific and time-dependent. While DNA had minimal effects on diffusion for most antibiotics, it significantly hindered the diffusion of, e.g., gentamicin and colistin at specific time points.

### 2.5 Antibiotic Diffusion through Mucoid Biofilms: Comparative Analysis with *In Vitro* Biofilm Models

A comparison using the setup in **Figure S5** of antibiotic diffusion through a 3-day-old mucoid biofilm (**Figures S6 and S7**), primarily composed of alginate, with other *in vitro* biofilm models revealed significant differences in diffusion kinetics, as shown in **Figure 2**. The diffusion of colistin and gentamicin was completely hindered at all time points, up to 20 h. For tobramycin, diffusion was negligible at 4 and 8 h; however, by 20 h, its diffusion reached levels comparable to those observed in the *in vitro* biofilm models, implying a significant antibiotic-matrix interaction until the matrix binding capacity was saturated. Ciprofloxacin, aztreonam, and meropenem exhibited a progressive increase in diffusion from 4 to 20 h through mucoid biofilms. Ciprofloxacin demonstrated a significantly lower diffusion at 4 h (p ≤ 0.001) through mucoid biofilms but reached equilibrium levels by 20 h. In the case of aztreonam, diffusion through the mucoid biofilm was significantly reduced by 4 and 8 h (p ≤ 0.001), while in meropenem, diffusion through the mucoid biofilm was significantly reduced by 8 and 20 h (p ≤ 0.05). While our models contained controlled concentrations of acetylated alginate and DNA, native mucoid biofilms include additional matrix components in addition to high levels of alginate and eDNA.^[43, 44]^ It is possible that these additional components or a different matrix structure can cause the observed significantly reduced antibiotic diffusion. This highlights the importance of matrix complexity in limiting drug access.

### 2.6 Role of Acetylation and DNA in Modulating Antibiotic Susceptibility

To assess the protective effects of alginate (acetylated and non-acetylated) with or without DNA on PAO1, bacterial susceptibility was evaluated using 2× the minimum inhibitory concentrations (MICs) based on EUCAST MIC breakpoints (**Table 1**). Results indicated that bacterial populations were reduced by ≥4 log after treatment with 2× MIC for 24 h compared with the control (p ≤ 0.05) for all models (**Figure 3**). However, interestingly, PAO1 exhibited increased tolerance to tobramycin, gentamicin, and ciprofloxacin in acetylated alginate biofilm models compared to non-acetylated alginate biofilm models. The opposite effect was observed for meropenem, aztreonam, and colistin, which were significantly more effective within an acetylated alginate matrix. The addition of DNA to the acetylated alginate biofilm model reduced tolerance, rendering PAO1 significantly more susceptible to gentamicin, ciprofloxacin, colistin, and meropenem (p ≤ 0.05).

**Figure 3.**
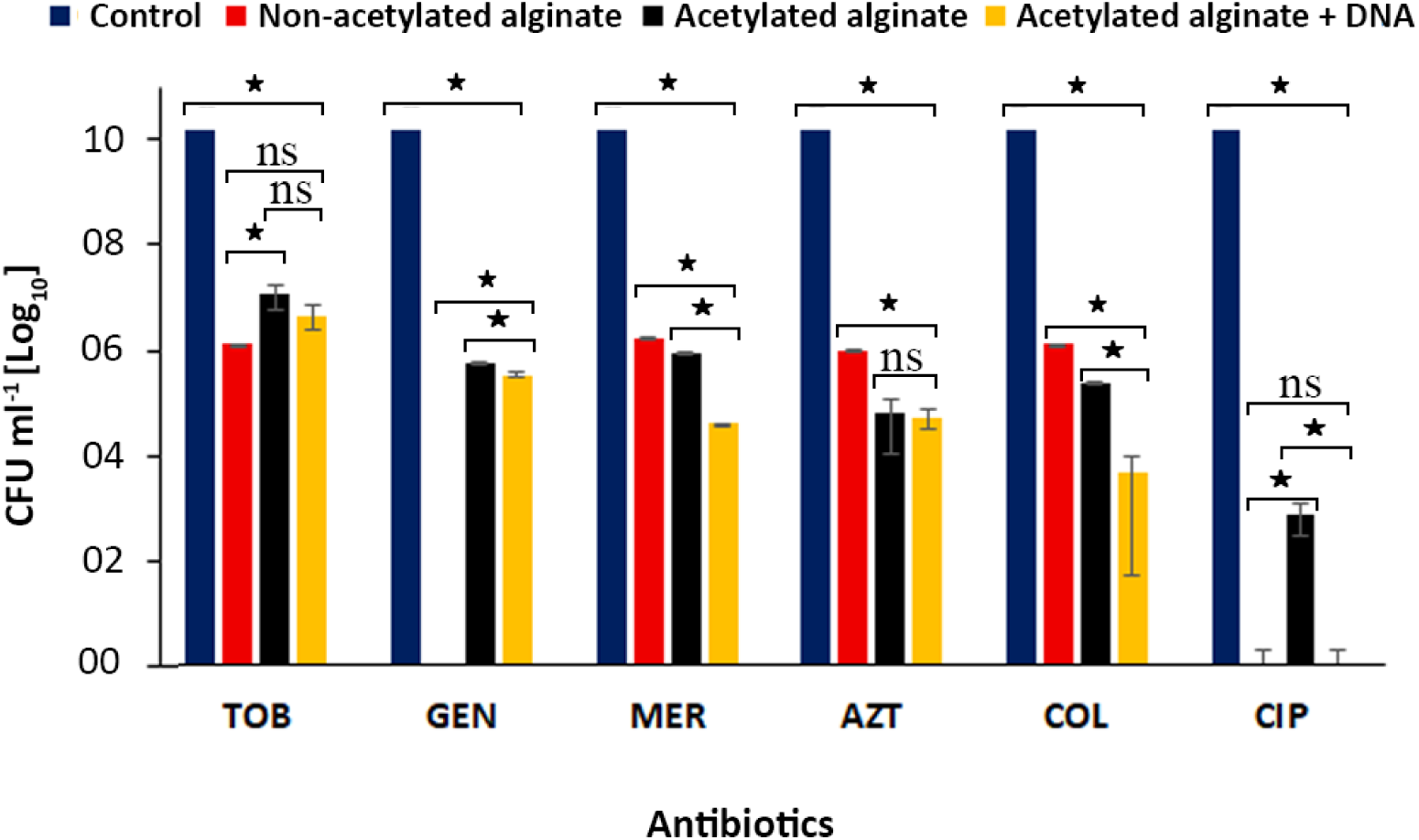
Chart representing the effects of different alginate-based encapsulation matrices on the susceptibility of PAO1. PAO1 was encapsulated in acetylated alginate and cross-linked with calcium and incubated for 1 h, after which it was exposed to treatment with 2x MIC [µg ml^−1^] (Table 1) of the selected antibiotics for 24 h at 37 °C. Surviving cells were counted by plating. * p ≤ 0.05, ns = non-significant.

**Table 1:** Minimum Inhibitory and Minimum Bactericidal Concentrations (MICs and MBCs) of selected antibiotics against PAO1. ECOFF = EUCAST epidemiological cut-off value.

|  | TOB | GEN | MER | AZT | COL | CIP |
| --- | --- | --- | --- | --- | --- | --- |
| | $\mu\text{g ml}^{-1}$ | | | | | |
| **MIC** | 1 | 0.06 | 0.25 | 1 | 0.25 | 1 |
| **MBC** | 4 | 1 | 1 | 2 | 0.5 | 16 |
| **MIC –ECOFF** | 2 | 8 | 2 | 16 | 4 | 16 |

### 2.7 Concentration-Dependent Antibiotic Killing of Encapsulated PAO1

The bactericidal activity of the selected antibiotics against encapsulated PAO1 was assessed by evaluating the relationship between increasing antibiotic concentration (µg ml^−1^) and bacterial viability, as demonstrated in **Figure 4**. Bacterial counts generally decreased to zero as antibiotic concentrations increased to 1000 µg ml^−1^, except for meropenem and aztreonam. Encapsulated PAO1 exhibited substantial tolerance to meropenem and aztreonam across all models, even at high antibiotic concentrations, with ≥ 99.9% eradication of bacterial cells requiring > 1024 µg ml^−1^ in all cases. The only exception was in non-acetylated alginate biofilm models, where complete eradication (≥99.9%) was achieved at 1024 µg ml^−1^ of meropenem.

**Figure 4:**
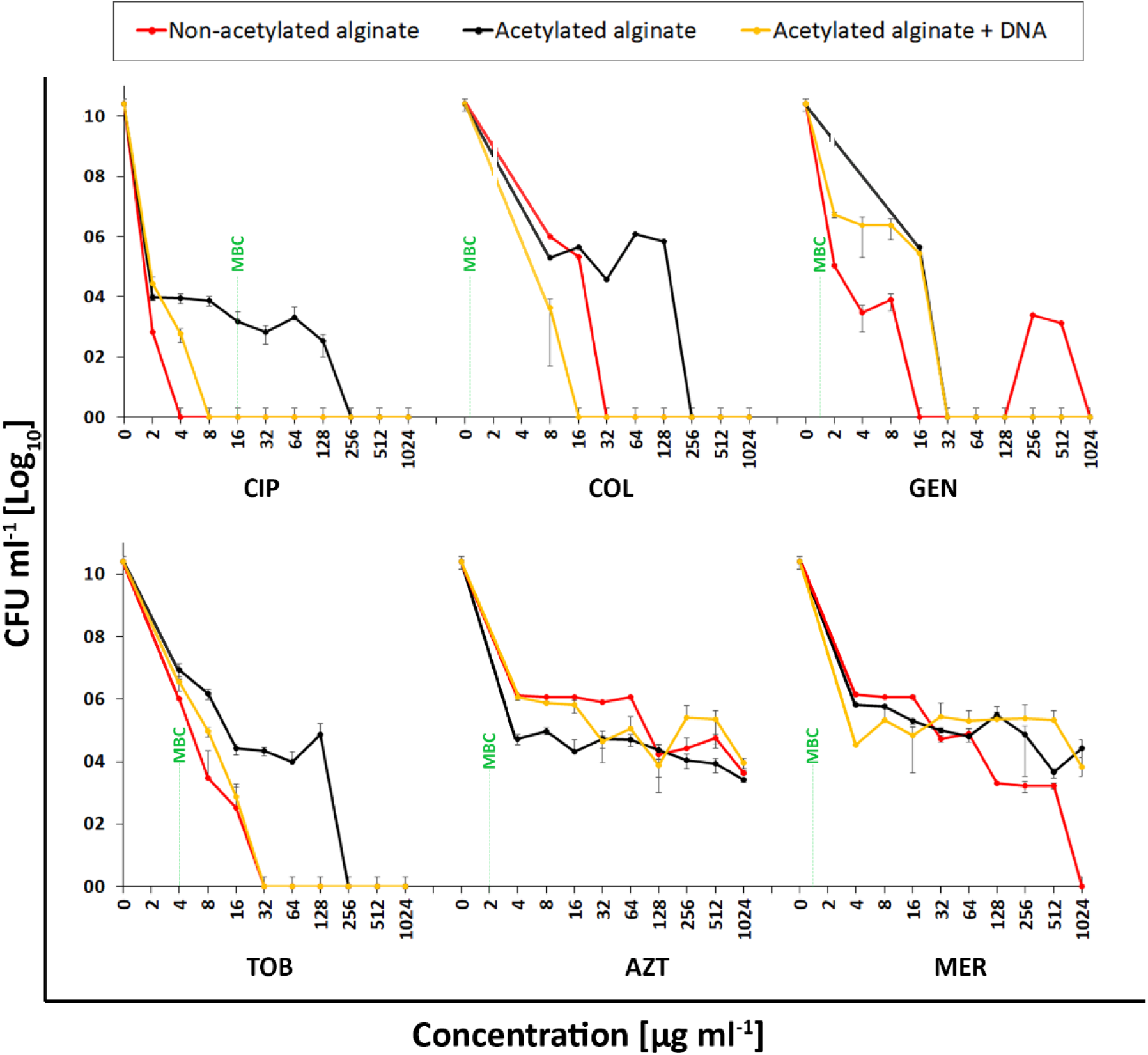
Concentration-dependent antibiotic killing of *P. aeruginosa* in *in vitro* biofilm models. PAO1 cells were encapsulated for 1 hour and subjected to antibiotic therapy for 24 h at various antibiotic concentrations. Surviving cells were counted by plating.

For tobramycin, ciprofloxacin, and colistin, alginate acetylation increased the antibiotic concentrations required to eradicate ≥ 99.9% of bacterial cells by more than 4-fold compared to their non-acetylated counterparts. However, adding DNA to the acetylated alginate biofilm model reverted this effect, reducing the concentration required to eradicate ≥ 99.9% bacterial cells by more than 3-fold.

### 2.8 Variations in Antibiotic Sensitivity of *P. aeruginosa* in Biofilm Models

Figure 5 compares the minimum biofilm inhibitory and eradication concentrations (MBIC and MBEC, respectively) with MICs (Table 1) for PAO1 encapsulated in the *in vitro* biofilm models. It revealed that encapsulating PAO1 in acetylated alginate, with or without DNA, yielded MBIC values more than three times the MIC in all cases. Notably, colistin and aztreonam MBICs were up to 1000 times the MIC (0.25 and 1µg ml^−1^, respectively) in acetylated alginate biofilm models.

**Figure 5:**
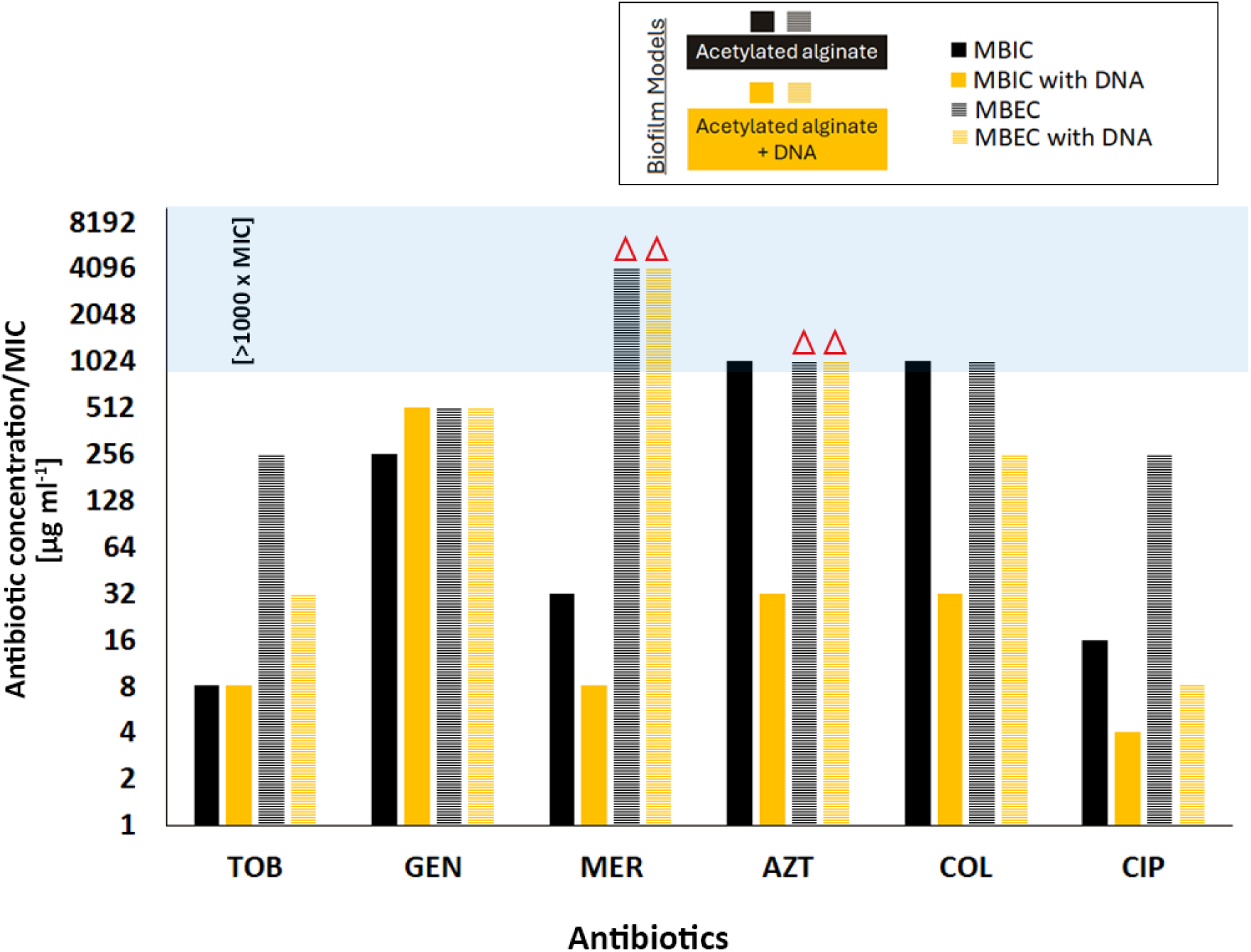
The chart presents MBIC and MBEC values (expressed as multiples of MIC for six antibiotics—Tobramycin (TOB), Gentamicin (GEN), Colistin (COL), Ciprofloxacin (CIP), Azithromycin (AZT), and Meropenem (MER)—against *P. aeruginosa* biofilms. The biofilm assays were conducted using acetylated alginate biofilm models, with and without DNA supplementation, to assess their impact on antibiotic tolerance. The red triangles represent eradication concentrations exceeding the measured range.

The minimum biofilm eradication concentrations (MBEC) exceeded 8x the MIC across all conditions, with aztreonam and meropenem requiring more than 1024 times the MIC (1 and 0.25 µg ml^−1^, respectively) for eradication. In evaluating the role of DNA in increased antibiotic tolerance, we observed that the addition of DNA to acetylated alginate neither increased nor reduced the eradication concentrations of gentamicin, aztreonam, or meropenem but increased the susceptibility of PA to tobramycin, ciprofloxacin, and colistin.

## 3. Discussion

The mucoid phenotype of PA is a leading cause of chronic lung infections in CF patients, characterized by overproduction of the EPS alginate. ^[38, 45, 46]^ A key chemical feature of alginate produced by mucoid PA is the presence of O-acetyl modifications, in which acetyl groups are covalently attached to hydroxyl groups of the M units in the polymer backbone.^[37, 39]^ This modification is notably absent in commercially available brown algae-derived alginate, which is commonly used in *in vitro* biofilm models.^[40]^ However, acetylation is hypothesized to be an adaptive mechanism that enhances biofilm structural integrity and confers additional physiological advantages, particularly within the CF lung environment.^[37]^ Acetylation reduces the charge density and hydrophilicity of the alginate and can shield the carboxyl groups participating in divalent (Ca^2+^) ion complexation and crosslinking. ^[16, 47, 48]^ A direct impact of acetylation on the diffusion and activity of antibiotics based on charge, size, and polarity could be expected, which makes the use of seaweed-derived alginate for biofilm models unsuitable.

The encapsulation approach employed in this study offers a reproducible model for investigating biofilm-associated infections, with potential applications in high-throughput testing of novel antimicrobial strategies. Encapsulation of PAO1 in both acetylated and non-acetylated alginate revealed distinct microcolony formation within 24 hours. These findings align with previous studies demonstrating that alginate-encapsulated PA formed structured biofilms in hydrogel-based matrices, mimicking *in vivo* conditions more accurately than traditional static biofilms. ^[49, 50]^

Biofilms are known to possess unique structural and biochemical properties that can restrict the diffusion of antibiotics, complicating therapeutic interventions.^[51, 52]^ Several studies report that alginate in biofilms impeded the diffusion of certain antibiotics like aminoglycosides, thereby contributing to antibiotic tolerance.^[11, 32, 43, 53, 54]^ However, other studies report little or no impairment of aminoglycoside diffusion through alginate matrices. ^[52, 55]^ To understand how the diffusion of antibiotics is suppressed, we created 3D biofilm models incorporating two key components: DNA and alginate. Additionally, we introduced acetylated alginate, representative of the alginate found in PA mucoid biofilms, and compared antibiotic diffusion through these models with that in native mucoid biofilms. Our results indicated that all three *in vitro* biofilm matrices (non-acetylated, acetylated, and acetylated + DNA) allowed antibiotic diffusion, with concentrations exceeding 80% after 8 hours, except for gentamicin, which was slightly lower. Importantly, our models showed that neither DNA addition nor alginate acetylation meaningfully affected antibiotic diffusion. The observed variations in diffusion rates were not statistically significant, and diffusion reached equilibrium levels within less than 20 hours. Our results, therefore, strongly suggest that reduced diffusion of antibiotics may not be a major reason for the observed increased tolerance.

The role of eDNA in enhancing antibiotic tolerance and binding antibiotics has been previously discussed. ^[56–58]^ In line with previous results on biofilms, we found that incorporating DNA in biofilm models notably reduced the diffusion of gentamicin and ciprofloxacin through the alginate matrix. Previous studies have suggested that negatively charged eDNA could bind cationic antibiotics, such as aminoglycosides, through electrostatic interactions. ^[21, 59]^ While a reduction in gentamicin diffusion supports this notion, tobramycin — another aminoglycoside with a similar charge — exhibited comparable diffusion rates between models with and without DNA. This observation implied that electrostatic interactions alone do not fully account for the modulation of antibiotic diffusion in the presence of eDNA. Moreover, molecular size appeared not to be the determining factor, as gentamicin (Mw 477.6 g mol^−1^) is smaller than tobramycin (Mw 565.6 g mol^−1^), yet its diffusion was more impeded. Notably, gentamicin is also less hydrophilic than tobramycin (logP −3.1 vs. −5.8, respectively), which may partially explain the observed differences in diffusion rate. The greater hydrophilicity of tobramycin likely facilitates its mobility within the more hydrophilic alginate–DNA matrix, indicating that hydrophobicity, in addition to charge, determines antibiotic transport in biofilm environments. Given that eDNA is known to contribute to biofilm acidification and ciprofloxacin exhibits higher solubility in acidic conditions, ^[59]^ an increase in the diffusion of fluoroquinolones would be expected. However, our results contradicted this hypothesis, demonstrating a significant reduction in ciprofloxacin (Mw 331.34 g mol^−1^) diffusion. This also suggested that factors beyond charge interactions and molecular size were responsible for the observed diffusion patterns.

An important consideration in interpreting our findings lies in the nature of the DNA (from Calf thymus) used. This preparation was selected as a standardized and reproducible model of bacterial eDNA due to its biochemical and structural similarities to bacterial eDNA. Both molecules are polyanionic, phosphate-rich macromolecules that interact with divalent cations (Mg²⁺, Ca²⁺), form electrostatic complexes with polysaccharides such as alginate and Psl,^[60, 61]^ and contribute to matrix viscoelasticity and charge distribution.^[60–62]^ These shared physicochemical properties justify the use of calf thymus DNA as a model substrate to evaluate DNA-mediated effects on antibiotic diffusion and biofilm structure. Nonetheless, we recognize that native eDNA produced within PA biofilms could differ markedly in its sequence composition, strand length distribution, conformational state, and chemical modifications.^[63, 64]^ Emerging evidence has also suggested that noncanonical DNA structures, such as G-quadruplex (G4) and Z-DNA, could exist within biofilm matrices and could exhibit unique functional and mechanical properties distinct from unstructured B-DNA (the form present in Calf thymus DNA). ^[61, 65, 66]^

Further investigation into the physical and chemical dynamics within the biofilm matrix is warranted to elucidate the mechanisms hindering some antibiotic diffusion in the presence of DNA. Additional factors could be, e.g., hydrophobicity and hydrogen-binding ability, which could be relevant for acetylated alginate and eDNA, respectively.

In 3-day-old mucoid biofilms, primarily composed of bacterial alginate, our results revealed significant differences in diffusion dynamics. We observed that colistin and gentamicin were completely hindered from penetrating the mucoid biofilm at all time points, up to 20 hours. These results align with previous research, which suggested that aminoglycosides, such as gentamicin, as well as polymyxin (colistin), face substantial hindrance in biofilm environments, particularly in mucoid biofilms. Cationic antibiotics such as tobramycin, colistin, and gentamicin were expected to interact electrostatically with the negatively charged alginate, eDNA, and other components of the biofilms, potentially leading to reduced diffusion.^[32, 67–70]^ However, a comparison with our model biofilm matrices showed that the attractive charge interaction of antibiotics with the two main components of such biofilms (alginate and eDNA) could not fully explain the poor diffusion, at least within seaweed-derived acetylated alginate matrices used as a proxy for bacterial alginate. In native mucoid biofilms, the matrix complexity and molecular composition could amplify charge-based interactions, potentially altering antibiotic diffusion and retention. Another consideration may be the differences in polymer density between the native mucoid bacterial alginate and the modified sea-weed alginate. While it might be intuitive to assume that native mucoid alginate is more densely packed and crosslinked than purified seaweed-derived alginate, our previous data do not support this assumption.

Based on the results reported in our referenced publications,^[16, 36]^ the mesh size analysis indicates that the mucoid PA alginate was not more densely crosslinked than the acetylated synthetic alginate. Specifically, the acetylated alginate with a higher degree of acetylation (36% d.ac.) exhibited a mesh size of approximately 40 nm, whereas the mucoid PA alginate displayed the largest mesh size among the tested samples, sometimes even greater than that of the modified alginates. Because a larger mesh size corresponds to a less densely crosslinked network, these findings suggest that mucoid alginate is not more compact than chemically acetylated alginate and would not structurally impede the diffusion of antibiotics more than mucoid PA biofilm alginate. In summary, the slow and variable diffusion of antibiotics, such as tobramycin, ciprofloxacin, aztreonam, and meropenem, through native biofilms compared to our models emphasizes the need for alternative treatment strategies that can enhance antibiotic diffusion or circumvent the protective biofilm matrix but are not exclusively focused on charge interactions.

The contrast between our synthetic alginate–DNA models and native mucoid biofilms underscore the complexity of antibiotic diffusion within natural biofilm environments. While our model systems allowed controlled investigation of alginate acetylation and DNA effects, they do not fully replicate the molecular composition or organization of native biofilms. In vivo, PA produces additional EPS components, including Pel and Psl polysaccharides, as well as matrix-associated proteins, lipids, and extracellular vesicles. While a lesser part of the biomass, they could strongly modify matrix density, charge heterogeneity, and antibiotic binding. These components or their effect on the matrix structure likely account for the stronger diffusion barriers observed in native mucoid biofilms compared to our synthetic models, although differences in alginate and DNA molecular weight and DNA structure between our model and native biofilms could also play a role.

In evaluating the level of antibiotic tolerance, alginate encapsulation was found to greatly increase tolerance, as evidenced by the tremendously elevated antibiotic concentrations required for complete biofilm eradication. Our findings from the diffusion studies suggested that sequestration of antibiotics by alginate was unlikely to be the primary factor driving enhanced biofilm tolerance, as the increased tolerance was observed even when antibiotic concentrations in the matrix could reach saturation levels, despite that highly cationic antibiotics such as tobramycin, gentamicin, and colistin could be affected by the crosslinked alginate matrix and its acetylation. ^[71, 72]^ In addition, the diffusion of poorly soluble ciprofloxacin and high-molecular-weight colistin (Mw = 1155 g/mol) could still be influenced by the crosslinked and acetylated alginate network, yet this did not appear to fully account for the observed levels of tolerance.

If antibiotic binding were the primary mechanism, we would expect strongly reduced or blocked antibiotic diffusion. Yet our findings showed that after a few hours, the bacteria were exposed to equilibrium concentrations of antibiotics but exhibited tremendous tolerance even in model biofilm matrices. This suggested that beyond sequestration, other biofilm-specific survival mechanisms were involved. To further emphasize this point, PAO1 encapsulated in alginate models also showed extreme tolerance to aztreonam and meropenem despite their high solubility and low molecular weight (∼435 g mol^−1^ and ∼383 g mol^−1^, respectively), giving them the ability to penetrate biofilms better than cationic and poorly soluble antibiotics.

Furthermore, the MBIC values were substantially higher than MIC values in all models, highlighting the protective nature of both the alginate and alginate + DNA matrix, consistent with prior research emphasizing the drastic difference in antibiotic susceptibility between planktonic and biofilm-encased bacteria. ^[51, 54, 55, 73]^ MBEC values were dramatically elevated, with meropenem and aztreonam requiring concentrations exceeding 1024× MIC for effective eradication, reinforcing the challenge of treating biofilm-associated infections with conventional antibiotic regimens. ^[6]^ Our findings agree with previous studies suggesting that alginate protects PA in biofilms against aminoglycosides and ß-lactams, which was partly attributed to reduced metabolic activity due to low oxygen and nutrient gradient. ^[25, 52, 74–76]^ Other studies have indicated that other mechanisms, such as the structural integrity of the biofilm matrix or the presence of specific exopolysaccharides, contribute to the increased antibiotic tolerance in biofilms.^[77, 78]^ However, a recent study by Liang *et al.* ^[74]^ found that the overproduction of exopolysaccharides Pel and Psl did not significantly affect tolerance to tobramycin, ciprofloxacin, and meropenem. Given that our simplified alginate matrix leads to an enormous increase in tolerance without affecting antibiotic exposure, our results reinforce the idea that biofilm-specific survival mechanisms triggered by being embedded in a 3D matrix are part of the key drivers of tolerance in mucoid PA biofilms. They also demonstrate that these mechanisms are present even without other components of the biofilm matrix, such as Pel or Psl.

The results in Figure 4 demonstrated perhaps the most complex interplay between alginate acetylation, DNA, and antibiotic efficacy in biofilm-encased PA. Notably, while increasing antibiotic concentrations generally reduced bacterial viability, substantial tolerance persisted for meropenem and aztreonam across all models, with eradication requiring concentrations exceeding 1024 µg ml⁻¹ — an observation consistent with prior studies indicating ß-lactams tolerance in biofilm-associated PA infections.^[79, 80]^ Interestingly, non-acetylated alginate allowed complete eradication at 1024 µg ml⁻¹ of meropenem, suggesting that acetylation, known to increase matrix viscosity and alter charge interactions, ^[16, 48]^ plays a significant role in shielding bacteria from beta-lactam antibiotics. For tobramycin, ciprofloxacin, and colistin, the effect of alginate acetylation was particularly striking, increasing the required concentration for ≥ 99.9% bacterial eradication by over fourfold. Although no studies have been performed to understand the role of acetylation in PA tolerance, our previous study^[16]^ demonstrated that acetylation of alginate could influence alginate’s mechanical and structural properties, which could create a microenvironment that promotes bacterial stress responses, further enhancing tolerance. Specifically, mechanical testing, isothermal titration calorimetry (ITC), and MRI-based diffusion analyses demonstrated that acetylation of the alginate backbone reduces the negative charge density and Ca²⁺ affinity of the alginate, leading to softer, more hydrated networks that diminish binding to positively charged aminoglycosides and facilitate their diffusion, thereby decreasing apparent binding. ^[16, 36]^

While the DNA-containing biofilm models initially exhibited a reduction in CFU ml^−1^ upon antibiotic treatment with aminoglycosides, ciprofloxacin, and colistin (Figure 4), which oddly correlated well with the antibiotics expected to show the strongest (electrostatic) interactions with DNA, this reduction was not significantly more pronounced than that observed in non-DNA matrices. Interestingly, adding DNA even reduced MBEC values for tobramycin, ciprofloxacin, and colistin (Figures 4 **and 5**). Adding DNA did not enhance susceptibility to meropenem and aztreonam, even though their diffusion was increased compared to alginate alone. These findings suggested that eDNA alone might not increase the antibiotic persistence of PA in biofilms. On the contrary, it could even weaken the protective mechanism provided by acetylated alginate. That DNA does not inherently confer increased tolerance to cationic or other tested antibiotics raises questions about the actual contribution of eDNA to biofilm resilience.

This apparent contradiction may be explained by several factors. According to literature, high levels of eDNA can substantially alter the physicochemical environment by depleting local cell-cations to an extent that compromises membrane stability of the cells, potentially increasing permeability to antibiotics rather than reducing it. ^[29, 81, 82]^ Furthermore, the binding of protons and ions by phosphate groups in DNA networks can locally acidify the matrix. Acidification may promote the uptake or potency of certain antibiotics whose activity is pH-dependent, thereby contributing to the observed reduction in tolerance. ^[57, 83, 84]^

Another plausible mechanism involves charge redistribution. In DNA-rich matrices, the strong negative charge of eDNA may attract cationic antibiotics more efficiently into the biofilm interior, effectively increasing their local concentration near bacterial cells.^[29, 30, 85]^ While this mechanism is opposite to the expected sequestration model,^[29, 56, 86]^ it aligns with findings that electrostatic gradients within biofilms can direct molecular diffusion in unintuitive ways.^[86, 87]^

Traditionally, eDNA has been viewed as an anionic, critical matrix component that fortifies biofilm structural integrity, promotes bacterial adhesion, and sequesters antibiotics, thereby limiting their efficacy.^[88–90]^ However, our results suggested that the role of eDNA in the antibiotic tolerance of biofilms is more nuanced and may not align with the conventional view that eDNA traps antibiotics like aminoglycosides through electrostatic interactions, reducing their bioavailability.^[88, 90]^

Finally, the comparison between our model systems and natural mucoid biofilms highlighted an essential point: although they capture PA resilience, our model systems do not replicate the additional molecular penetration barrier observed in *in vivo* biofilms. As shown in Figure 2, real biofilms introduce diffusion limitations that may be due to additional matrix components or a denser matrix structure. Our models, however, demonstrated that even in the absence of a significant reduction in antibiotic diffusion, bacterial survival remained alarmingly high. The disconnect between antibiotic efficacy and diffusion highlighted the role of biological rather than purely physical mechanisms in driving biofilm-associated tolerance. Some biological mechanisms may include efflux pump expression, proteins, lipids, and extracellular vesicles in PA biofilms. ^[7, 45, 91–93]^ Hence, antibiotic failure in chronic infections is not merely a problem of access but also bacterial physiological state and biofilm physiology. Ultimately, these findings advocate for a shift in focus from overcoming diffusion barriers to targeting the bacterial adaptations triggered by encapsulation that sustain biofilm antibiotic tolerance. Integrating transcriptomic or proteomic analyses to determine how matrix-derived mechanical cues modulate bacterial physiology and antibiotic tolerance could provide valuable insight into this phenomenon.

In parallel, improving therapeutic efficacy in patients will also require strategies that enhance local availability of antibiotics by optimizing administration routes, formulation design, and how long the drug remains active to maintain effective concentrations against persistent bacterial populations.

While PAO1 served as a standardized model for this study, clinical CF isolates often display distinct phenotypes, including differences in antibiotic susceptibility. Future studies will therefore assess whether mucoid and non-mucoid clinical isolates exhibit similar antibiotic tolerance patterns within acetylated alginate ± DNA matrices. Such comparative studies will be essential to validate the broader clinical relevance of our findings and determine whether the mechanisms identified here extend to biofilms formed by patient-derived PA strains.

## 4. Conclusion

We conclude that neither alginate alone nor alginate with DNA significantly impedes antibiotic diffusion, in contrast to mucoid biofilms that strongly suppress antibiotic diffusion. Instead, we conclude that the barrier effect observed in mucoid *P. aeruginosa* biofilms likely arises from matrix properties such as high density and compositional complexity. However, it remains possible that in native biofilms, direct interactions between antibiotics and the high molecular weight bacterial alginate and bacterial eDNA play a dominant role in diffusion retardation, an effect that may not be fully captured by the chemically modified alginate and commercially available DNA used in our model. Additional biofilm matrix components, including proteins, lipids, and extracellular vesicles, may contribute to the overall increased antibiotic tolerance by influencing antibiotic sequestration, neutralization, or altered microenvironmental conditions.

We demonstrated that embedding in printed alginate hydrogels massively increased PAO1 antibiotic tolerance despite the model biofilm matrix’s negligible influence on antibiotic availability. Interestingly, acetylation, in most cases, dramatically decreased antibiotic susceptibility, indicating that acetylation is beneficial for antibiotic tolerance without reducing antibiotic diffusion, but the inclusion of DNA negates this effect. Our findings significantly strengthen the hypothesis that biofilm-associated tolerance mechanisms other than impaired antibiotic diffusion play a crucial role and are triggered by encapsulation in a biofilm-like environment that can be printed. These may include metabolic dormancy, adaptive stress responses, efflux pump activation, and genetic adaptation, all of which can enhance bacterial survival in the presence of biofilm-penetrating antibiotics. Our results were obtained under the constraints of a simplified, biofilm-like model based on seaweed-derived alginate and cannot directly identify the underlying biological pathways. Given that tolerance in our study is triggered by encapsulation in fully synthetic matrices, it would be of highest interest to investigate these cellular responses with respect to a physicochemical environment modified both synthetically and with matrix components extracted from mucoid biofilms. Furthermore, studying the synergistic roles of other extracellular biofilm components with acetylated alginate and DNA in affecting antibiotic transport, elucidating the molecular mechanisms underlying these interactions, and exploring how bacteria-matrix interactions trigger antibiotic tolerance will be critical to confirm the triggering and origin of specific adaptive mechanisms directly. This could lead to combination therapies that negate these matrix effects to enhance bacterial eradication in biofilm-associated infections.

## 5. Experimental Section/Methods

### 5.1 Bacterial strains, growth conditions, and antibiotics

The bacterial strains used in this study are listed in Table S1. Unless otherwise noted, frozen bacterial strains were thawed and grown overnight at 37 °C in Tryptic Soy broth (TSB). Stock solutions for Tobramycin sulfate, Gentamicin, Ciprofloxacin, Aztreonam, Meropenem, and Colistin were prepared and further diluted in Mueller Hinton Broth (MHB) – pH 7.4 to the desired concentrations for further antibiotic studies.

### 5.2 Acetylation of Seaweed Alginate

As described in ^[16]^, alginate beads (1 wt%) were prepared and crosslinked for 24 h in 0.1 M CaCl_2_ after which solvent was exchanged with pyridine at 0, 3, and 24 hours. Acetylation was performed by adding Ac_2_O (3.5 ml) in pyridine (96.5 ml). The reaction was stirred at room temperature for 24 h. Beads were filtered, washed twice with acetone and deionized water (100 ml per wash, with 10 min stirring between steps), and dissolved overnight in 50 ml of 0.2 M EDTA (pH 7.4). The solution was dialyzed (MWCO 3.5 kDa, Sigma-Aldrich) against 0.9% NaCl for 48 h, followed by deionized water for another 48 h. After dialysis, it was concentrated to half the volume using a rotary evaporator and lyophilized (Christ Gamma 2–20, Osterode am Harz, Germany). The final product, a white, cotton-like material, was stored at −20 °C under an N₂ atmosphere. The degree of acetylation was determined by ^1^H-NMR after enzymatic digestion (0.1 U alginate lyase, pH 6.5, 10 mg ml^−1^ alginate, 38 °C, 24 h). ^[94]^

### 5.3 Bioink Preparation

Bioink stock solution of 1.5 wt% was prepared by dissolving 150 mg of acetylated alginate or non-acetylated alginate overnight in 8 ml of sterile 0.9 wt% NaCl solution. Subsequently, the solvent was added to the mixture to reach the final volume of 10 ml. For bioink containing bacteria, the stock solution was diluted with 0.9 wt% NaCl, TSB (pH 7.4), and PAO1 to obtain a 0.5 wt% alginate solution as bioink with a defined bacterial concentration (0.01 OD_600_). For bioink containing DNA and bacteria (Figure 6), the stock solution was mixed with 0.9 wt% NaCl, TSB, DNA, and PAO1 to obtain a 0.5 wt% alginate and 0.1 wt% DNA solution as ink with a defined bacteria concentration (0.01 OD_600_). ^[95]^

**Figure 6:**
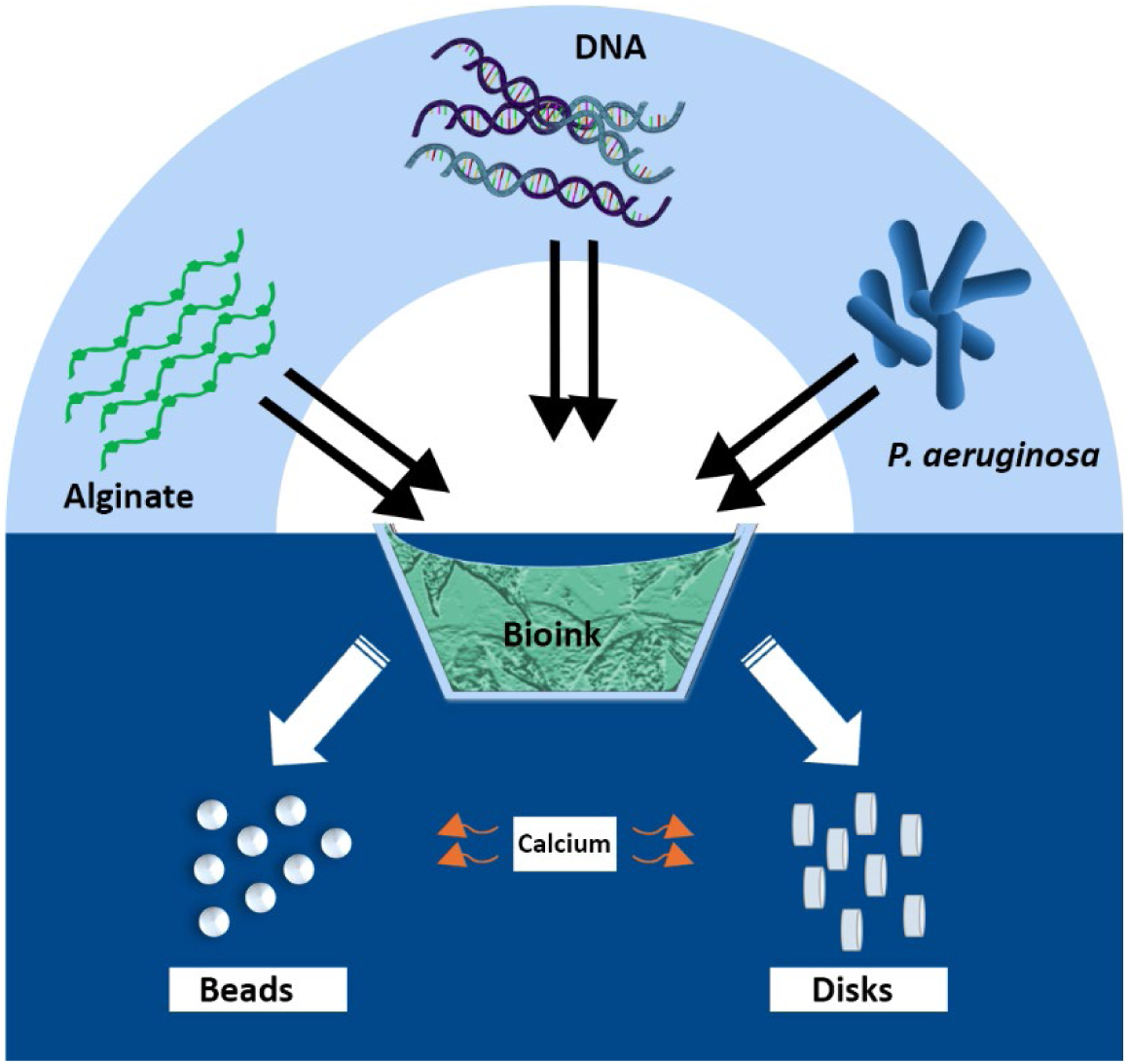
Graphical representation of 3D biofilm model fabrication. Bioink was prepared by mixing acetylated or non-acetylated alginate with PAO1, with or without DNA, and cast as disks or printed as beads, crosslinked with CaCl_2_ for 1 h.

### 5.4 Preparation of Alginate and Biofilm Models

To produce disk models, using a modified method,^[95]^ CaCl_2_-agarose molds were prepared by dissolving 2 g of agarose powder (Sigma) and 1.47 g of CaCl_2_ in 100 ml of deionized water, followed by autoclaving. The agarose was gelled to a height of 3 mm and punctured with 7 mm diameter punches. Approximately 150 µl of the respective bioinks containing alginate, PAO1, and/or DNA (see Section 5.3) were pipetted into the molds and allowed to crosslink for 1 h. To produce bead models, bioink (see Section 5.3) was loaded into a cartridge for a pneumatic pump, inserted into an EMD printhead (Cellink, BICO, Sweden), and mounted on a BioX Gen3 (Cellink, BICO, Sweden). The pressure applied to the formulation for inkjet printing ranged from 45 to 90 kPa, depending on the type of Alginate. The printing parameters for open and cycle time and writing speed were set to 1 and 150 ms and 1.5 mm s^−1^, respectively. The formulation was jetted into the wells of a 48-well plate containing 800 μl of a sterile, filtered 350 mM CaCl_2_ solution per well. The produced beads were left to crosslink in the solution for 1 h. The disks (∼7 mm) and beads (∼600 µm) from both methods were then carefully removed, washed three times with 0.9 wt% NaCl, and transferred to TSB for subsequent experiments.

### 5.5 Microscopy Analysis

To analyze the structure and viability of bacterial aggregates, alginate-encapsulated PAO1 was cultured in TSB for 8, 24, and 48 h. The biofilms were stained with LIVE/DEAD™ BacLight™ Bacterial Viability Kit (Thermo Fischer Scientific) and observed with a confocal laser scanning microscope (Leica SP8) in fluorescence mode using optimal excitation wavelength/emission wavelength (Ex/Em) of 485⁄498 nm for SYTO9 and Ex/Em of 535⁄617 nm for propidium iodide.

Samples for cryo-scanning electron microscopy (cryo-SEM) were high-pressure frozen using the Leica EM HPM100. The frozen samples were automatically discharged into a liquid nitrogen-filled dewar for preservation before being transferred to the Leica EM ACE900 freeze-fracture system under cryogenic conditions. Freeze-fracturing was performed at −120 °C using a cryogenic knife, followed by sublimation at −110 °C under a vacuum of 2.6 × 10^−6^ mbar for 20 s to enhance structural visibility. The samples were then coated with a 4-nm platinum layer at a 45° angle and a 4 nm carbon layer at a 90° angle through e-beam physical vapor deposition to improve contrast and minimize beam damage during imaging. Cryo-SEM imaging was performed using the Apreo VS SEM (Thermo Scientific, The Netherlands) at an operating voltage of 1.0 kV and a current of 0.10 nA in a high vacuum mode. Backscattered and secondary electrons were detected via the in-lens T1 and T2 detectors, respectively. Throughout the imaging process, the SEM stage temperature was maintained between −110 °C and −120 °C, depending on the chamber pressure to avoid sublimation and condensation, as observed *in situ*.

### 5.6 Antibiotic Diffusion Assay

#### 5.6.1 Biofilm Models and Controls

The assay was performed on native mucoid biofilms and three biofilm models: acetylated alginate, non-acetylated alginate, and acetylated alginate with DNA. 2 wt% agarose disks (∼7 mm diameter) were used as controls.

#### 5.6.2 Native Mucoid Biofilm Growth

Mucoid PA biofilms were grown with slight modifications. ^[43]^ A mucoid PA culture (OD₆₀₀ = 0.1) was pipetted onto agarose disks in Petri dishes and incubated for 1 h to allow bacterial attachment. Excess bacteria were removed by washing three times with 1 ml TSB. Subsequently, 50 µl of TSB was added to 145 mm Petri dishes containing agarose disks, ensuring the surface remained exposed to air while receiving nutrients from below. Bacteria-loaded agarose disks were incubated for 3 days at 37 °C for biofilm development on the surface of the disks.

#### 5.6.3 Antibiotic Diffusion Assessment

To evaluate antibiotic diffusion through biofilm models (non-acetylated alginate, acetylated, and acetylated alginate with DNA), one agarose disk was placed on top of all the models, forming a 2-stack assembly (Figure S4). As a control, agarose disks were placed on another agarose disk with or without DNA, forming a 2-stack assembly. For native mucoid PA biofilms of ∼0.25 mm thickness, an agarose disk was placed on the disk containing biofilms, forming a 2-stack assembly. An additional agarose disk was added to this stack to prevent the spread of mucoid PA onto the topmost disk. As a control, 3 agarose disks were used to form a 3-stack assembly. ^[43]^

All stacks were transferred to antibiotic-containing MHA plates with the following concentrations tobramycin (50 µg ml^−1^) gentamicin (50 µg ml^−1^), ciprofloxacin (5 µg ml^−1^), aztreonam (30 µg ml^−1^), meropenem (10 µg ml^−1^), and colistin (8 µg ml^−1^), chosen as concentrations that should kill *E. coli* (EUCAST). At designated time points, the topmost agarose disks were removed and placed onto MHA pre-inoculated with *E. coli* (OD₆₀₀ = 0.1). After 24 h incubation at 37 °C, the diameter of the inhibition zones was measured. Percent diffusion was determined using the formula:

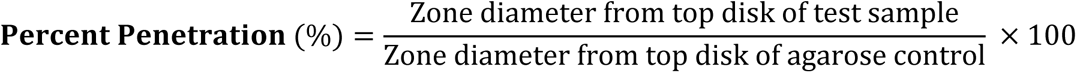

All experiments were conducted in triplicate.

### 5.7 Antimicrobial Susceptibility Assay

Minimum inhibitory concentrations (MICs) were determined using the broth microdilution method as recommended by the EUCAST. ^[96]^ Prepared PAO1 suspensions were added to equal volumes of Muller-Hinton Broth (MHB) containing two-fold dilution concentrations of each antibiotic in a sterile well-plate to get the final inoculum size of 5 × 10^5^ CFU ml^−1^. Pure PAO1 suspension was used as a positive control, and MHB as a negative control. The plates were incubated at 37 °C for 16 h and MICs were recorded as the lowest concentrations with no visible growth. To determine the minimum bactericidal concentration (MBC), 100 µl of the bacterial suspensions from plates with no visible growth were spread onto Muller-Hinton Agar (MHA) to observe viability after overnight incubation at 37 °C. The MBCs were recorded as the lowest concentrations of antibiotics that reduced the viability of the initial bacterial inoculum by ≥ 99.9%. All experiments were done in triplicate.

### 5.8 Biofilm Inhibition and Eradication Assay

*P. aeruginosa* PAO1 encapsulated in alginate disks (at an OD_600_ of 0.01) were incubated in TSB, pH 7.4, at 37 °C for 1 h to facilitate initial bacterial adaptation within the matrix. The MBIC assay was conducted by transferring the alginate-encapsulated PAO1 disks into a 96-well microtiter plate containing serially diluted antibiotics in MHB. The plates were then incubated at 37 °C for 24 h. Minimum Biofilm Inhibitory Concentration (MBIC) was determined by measuring the optical density at 600 nm (OD_600_) and visually inspecting turbidity. The MBIC was defined as the lowest antibiotic concentration at which an increase in OD_600_ could not be detected. For the Minimum Biofilm Eradication Concentration (MBEC) assay, after 24 h of antibiotic exposure, the disks were dissolved using a solution of 0.05 M Na_2_CO_3_ and 0.02 M citric acid. ^[97]^ The resulting suspension was diluted to a final volume of 1 ml with PBS. Bacterial regrowth was evaluated by plating 100 µl of each sample on three TSA plates (total plated volume = 300 µl) and incubating them overnight. This corresponds to a detection limit of approximately 3 CFU/mL, calculated based on the total sample and plated volume. The MBEC was determined as the lowest concentration at which no CFUs could be counted. All experiments were performed in triplicate.

### 5.9 Statistical analysis

All experiments were performed in biological and technical triplicate unless specified otherwise. Results are reported as mean ± standard deviation and were evaluated with one-way ANOVA With Tukey’s *post hoc* tests to compare means within and between groups.

## Author contributions

Goodness Osondu-Chuka: writing the original draft, primary investigation, analysis, and reporting. Stephan Schandl: review and editing, investigation, and analysis. Andrea Scheberl: formal analysis. Olivier Guillaume: review and editing, funding acquisition, project administration. Aleksandr Ovsianikov: review and editing, supervision. Erik Reimhult: writing, review, and editing, supervision, formal analysis, funding acquisition, project administration.

## Acknowledgements

This project was funded by FWF-Stand Alone “BREATH” Project with project number P33226.

## Data availability

The authors confirm that the data supporting the findings of this study are available within the article and its supplementary information.

## Code availability

This study did not generate any code

## Supplementary Information

Supporting Information is available from the npj biofilms and microbiomes library or from the author

