## Supporting Information for "Beyond the Matrix: Rethinking Antibiotic Tolerance in CF Biofilms Using 3D Models"

S. Schandl, O. Guillaume, A. Ovsianikov
Research Group 3D Printing and Biofabrication, TU Wien, 1060 Vienna, Austria

**Table S1**: List of bacteria used for all experiments

| | Bacteria | Strain |
|---|---|---|
| **1** | *Pseudomonas aeruginosa* | PAO-1 |
| **2** | *Pseudomonas aeruginosa* | Mucoid Strain – ATCC 39324 |
| **3** | *Escherichia coli* | DSM 1103 |

**Table S2.** List of media and reagents used

| | Medium/Reagent | Manufacturer |
|---|---|---|
| **1** | Tryptic Soy Broth | Sigma Aldrich - 22092 |
| **2** | LB Broth (Miller) | SIGMA Life Science L3522 |
| **3** | Pseudomonas Isolation Agar | Sigma Aldrich - 17208 |
| **4** | Calcium Chloride | ROTH – A119.1 |
| **5** | Muller Hinton Broth | Sigma Aldrich - 70192 |
| **6** | Agar | Sigma Aldrich - 05039 |
| **7** | Agarose (peqGOLD Universal agarose) | peqlAB 35-1020 |
| **8** | Tobramycin sulphate | Thermoscientific J62995.03 |
| **9** | Ciprofloxacin | CAS RN® 85721-33-1 |
| **10** | Meropenem | CAS RN® 119478-56-7 |
| **11** | Gentamicin sulfate | Sigma Aldrich G1264 |
| **12** | Deoxyribonucleic acid sodium salt from calf thymus | Sigma Aldrich D1501 |
| **13** | Colistin sulfate | CAS RN® 1264-72-8 |
| **14** | Aztreonam | CAS RN® 78110-38-0 |

### Cultivation and Extraction of Bacterial Alginate

Bacterial alginate was extracted from mucoid PA biofilms using a modified protocol.[1] Overnight cultures in tryptic soy broth (TSB) at 37 °C were harvested, washed three times in phosphate-buffered saline (PBS), and resuspended in TSB to $OD_{600}$ = 0.1. 200 µl was spread onto *Pseudomonas* isolation agar (PIA) plates and incubated at 37 °C for 72 h to develop biofilms. Biofilm biomass was scraped off, suspended in normal saline, and homogenized by magnetic stirring. The suspension was centrifuged at 25,000 × g for 30 min at 4 °C, and the supernatant was precipitated with ice-cold 2-propanol (1:1 v/v) overnight at 4 °C. The precipitate was recovered by centrifugation (15,000 × g, 30 min, 4°C) and washed thrice with ice-cold 2-propanol. To remove nucleic acids, the precipitate was resuspended in 50 mM Tris–HCl (pH 7.5) with 1 mM $CaCl_2$ and 2 mM $MgCl_2$, treated with DNase I and RNase A (15 mg $ml^{-1}$ each) at 37 °C for 6 h, followed by Proteinase K (5 mg $ml^{-1}$) digestion at 37 °C overnight. The solution was dialyzed (MWCO 14 kDa, Carl Roth) against 5 L Milli-Q water, lyophilized overnight, and obtained as an off-white, cotton-like powder. Alginate yield and uronic acid content were quantified using a carbazole assay with seaweed-derived alginate as a standard, with technical triplicates. Negative controls were included by treating samples with 0.0125 M NaOH.

### Nuclear magnetic resonance (NMR) spectroscopy and Gel permeation chromatography (GPC)

For $^1$H-NMR analysis, samples were enzymatically digested with alginate lyase according to the manufacturer's protocol for 24 h. Ethanol was used to precipitate the resulting digest, and the pellet was dried overnight at 60 °C. The dried material was subsequently dissolved in deuterated water ($D_2O$). The degree of acetylation was quantified following the method described by. [2] $^1$H-NMR spectra of alginate solutions were recorded on a BRUKER Avance DRX-400 FT-NMR spectrometer and analyzed using the MestReNova software by Mestrelab Research. To determine the number-average molecular weight ($M_n$) and polydispersity index (PDI) of alginate samples (2 mg $ml^{-1}$ in PBS), gel permeation chromatography (GPC) was performed using a Viscotek GPC system. The eluent consisted of PBS buffer (5 mM $NaH_2PO_4$, 100 mM NaCl, 0.05% $NaN_3$, pH 7.4) and was maintained at 30 °C with a flow rate of 1 ml $min^{-1}$.

As recently reported,[3] the synthetic acetylation does not degrade the molecular weight of the starting material and rather causes an increase due to the addition of the acetyl side group, as summarized in **Table S3**. Furthermore, the polydispersity index (PDI) decreases slightly due to dialysis purification and loss of small molecular weight fractions. $^{1}$H-NMR further confirmed the presence of the acetyl side group at a chemical shift of 2.0-2.2. While a similar degree of acetylation to the native mucoid alginate could be achieved, the acetylation pattern could not be controlled in the synthetic acetylation. Additionally, the digestion using an alginate lyase (polymannuronase) was drastically hindered by the presence of the acetyl side group. Native mucoid alginate showed the lowest resolution, which could be interpreted as the least digested alginate in this study.

**Table S3**. Molecular weight comparison obtained by GPC measurements. Synthetically acetylated alginate showed increasing molecular weight and lower polydispersity index (PDI) compared to the starting material. The isolated native mucoid alginate showed roughly three times the molecular weight of the seaweed-derived alginate used in this study.[3]

| **Alginate** | **$M_n$ (kDa)** | **PDI** |
|---|---|---|
| **0% degree of acetylation** | 126 | 2.7 |
| **36% degree of acetylation** | 147 | 2.3 |
| **Native mucoid** | 423 | 1.1 |

**Preparation and Scanning Electron Microscope Imaging of Alginate Hydrogel Samples**

The samples for scanning electron microscopy were first chemically fixed by immersing them in a 2.5% glutaraldehyde solution at 4 °C for 1 h. After fixation, the samples were rinsed twice with a 0.9 wt% NaCl solution to remove excess fixative. They were then dehydrated by gradually increasing the ethanol concentration in the following steps: 30%, 50%, 70%, 90%, 95%, and 100% ethanol, with each concentration incubated for 3 minutes at room temperature. Following dehydration, the samples were transferred to acetone as an intermediary solvent before starting the critical point drying (CPD) process with the Leica EM CPD030. After drying, the samples were sliced and coated with ~4 nm gold before imaging using a Apreo VS Scanning Electron Microscope (SEM) (Thermo Scientific, The Netherlands).

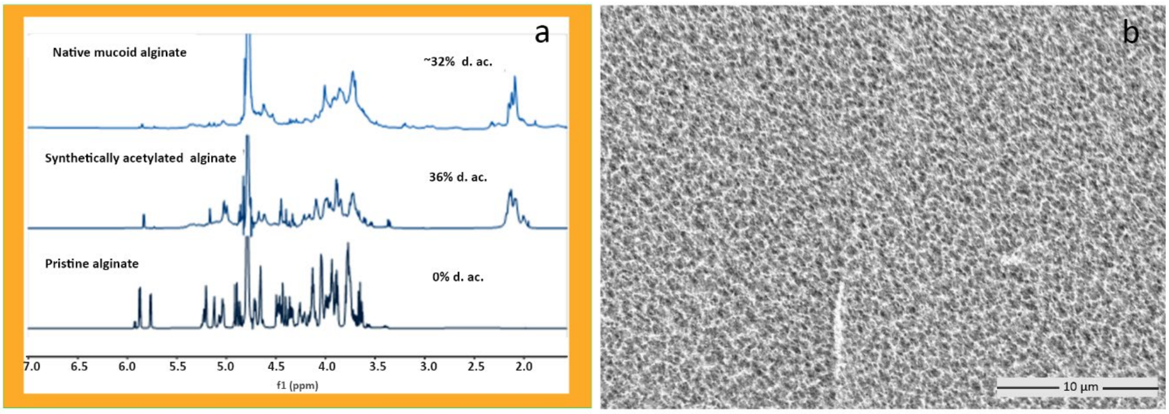


**Figure S1**: a) [1]H-NMR spectra of pristine alginate, its synthetically acetylated analog, and mucoid alginate isolated from a three-day-old *P. aeruginosa* biofilm. b) Cryo-SEM micrograph of acetylated alginate

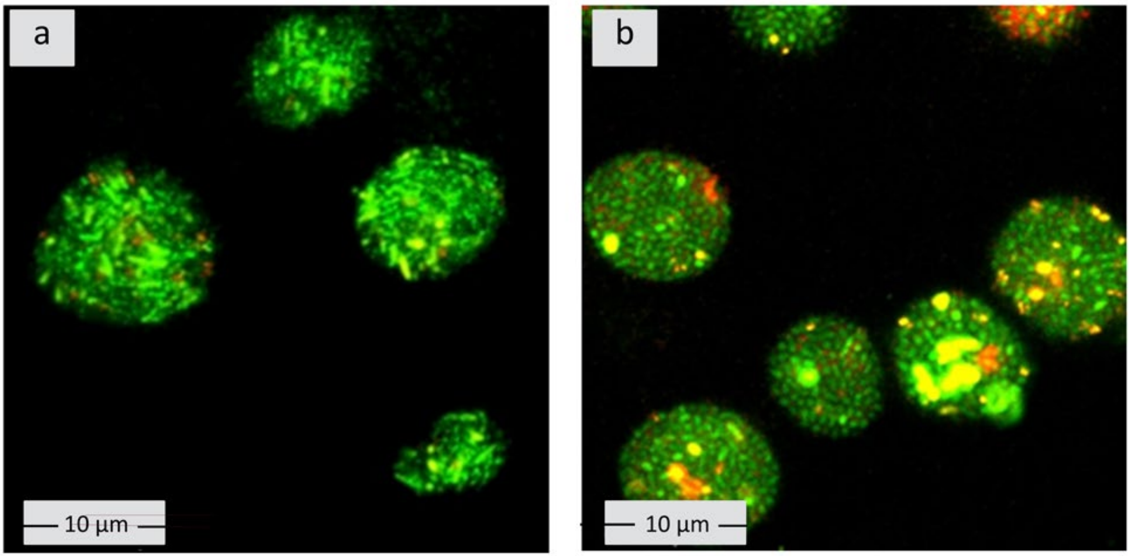


**Figure S2**. CLSM microscopy images of *P. aeruginosa* (PAO1) microcolonies in alginate biofilm models. Alginate-encapsulated PAO1 was cultured in TSB for 24 h and 48 h. The biofilms were stained with LIVE/DEAD™ BacLight™ Bacterial Viability Kit and observed with CLSM in fluorescence mode using optimal excitation wavelength/emission wavelength (Ex/Em) of 485⁄498 nm for SYTO9 and Ex/Em of 535⁄617 nm for propidium iodide. a) and b) PAO1 microcolonies at 24 h and 48 h, respectively.

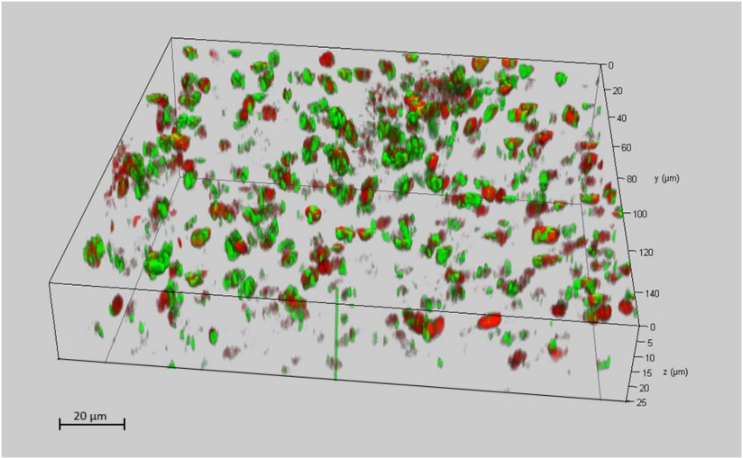


**Figure S3**. 3D CLSM image of 24 h PAO1 microcolonies in alginate biofilm model.

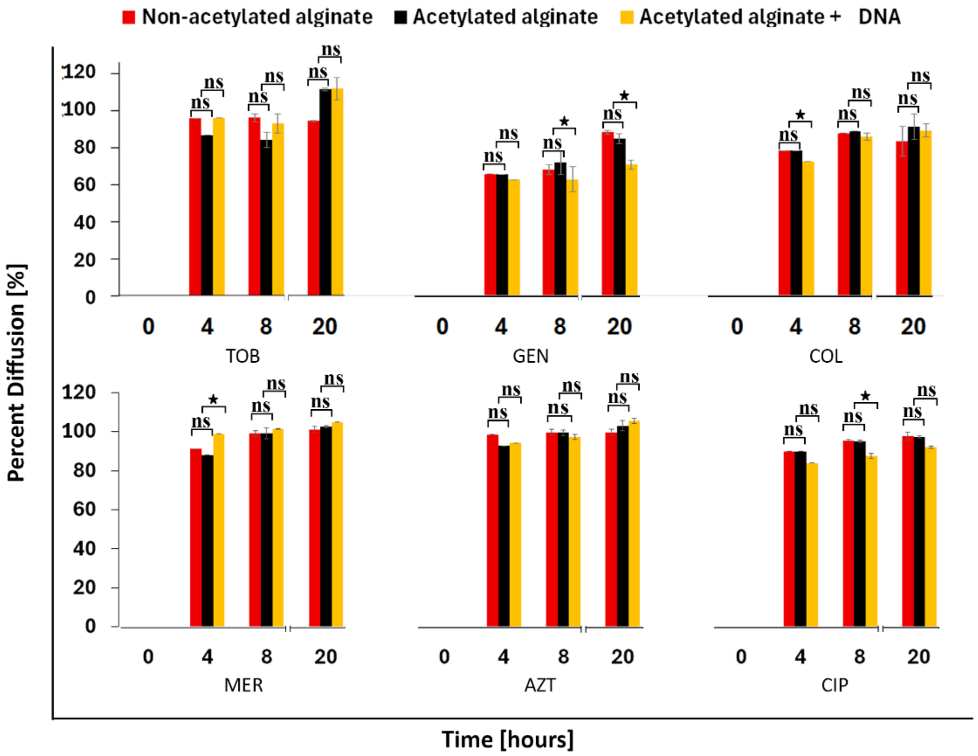


**Figure S 4**. The chart represents diffusion percentages of selected antibiotics through different 3D biofilm models and native biofilms at different time points. Percent diffusion is calculated relative to diffusion through an agar disk. One agarose disk was placed on top of each biofilm model, forming a 2-stack assembly. A 2-layer agarose disk stack was assembled for the reference measurements. Stacks were placed on MHA plates containing antibiotics (tobramycin, gentamicin, ciprofloxacin, aztreonam, meropenem, colistin) at concentrations effective against *E. coli*. At defined time points, the topmost agarose disk was transferred to *E.*

*coli*-inoculated MHA to assess antibiotic diffusion via inhibition zone measurement after 24 h at 37 °C. * $p \leq 0.05$, ns = non-significant.

**Table S4.** Physicochemical properties of antibiotics used in the manuscript were obtained from references [4–6].

| AB | Charge | pKa | Solubility | MW ($g.mol^{-1}$) | Ring Structure | AB Class | Relevance in CF Therapy | Method of Administration | Mode of action |
|---|---|---|---|---|---|---|---|---|---|
| **TOB** | Cationic | ~8.2 | Highly water-soluble | 467.52 | Aminocyclitol | Aminoglycoside | Yes | Intravenous, inhalation | Inhibition of protein synthesis |
| **GEN** | Cationic | ~8.2 | Highly water-soluble | 477.6 | Aminocyclitol | Aminoglycoside | Yes | Intravenous, inhalation | Inhibition of protein synthesis |
| **COL** | Cationic | NA | Partially water-soluble | 1,175.45 | Cyclic peptide | Polymyxin | Yes – last resort | Intravenous, inhalation | Disruption of bacterial cell membrane |
| **CIP** | Zwitterionic | ~4.04 | Partially water-soluble | 331.34 | Quinoline | Fluoroquinolone | Yes – not typically | Oral, intravenous | inhibits DNA gyrase |
| **MER** | Zwitterionic | NA | Partially water-soluble | 383.5 | β-lactam | Carbapenem | Yes | Intravenous | Inhibition of bacterial cell wall synthesis |
| **AZT** | Zwitterionic | NA | Partially water-soluble | 435.4 | β-lactam | Monobactam | Yes | Intravenous, inhalation | Inhibition of bacterial cell wall synthesis |

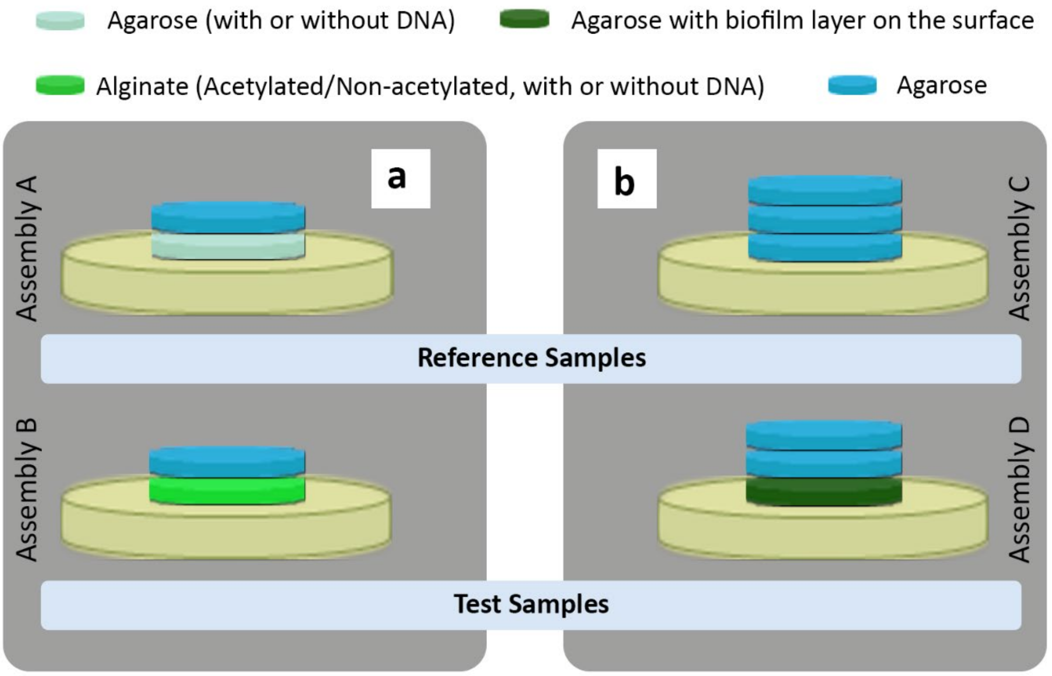


**Figure S5**. Schematic representation of the antibiotic diffusion assay. To evaluate antibiotic diffusion through biofilm models (acetylated, non-acetylated, and acetylated alginate with DNA) and mucoid P. aeruginosa biofilms, one agarose disk (~ 1,6 mm height, ~ 7 mm diameter) was placed on each biofilm model (~ 1,6 mm height, ~ 7 mm diameter), forming a 2-stacked assembly (Assembly A). For stacks with mucoid biofilms, an extra agarose disc was added to prevent the spreading of P. aeruginosa onto the topmost disc (Assembly C). These stacks were placed on MHA with the following antibiotic concentrations: 50 µg $ml^{-1}$ tobramycin sulfate, 50 µg $ml^{-1}$ gentamicin, 5 µg $ml^{-1}$ ciprofloxacin, 30 µg $ml^{-1}$ aztreonam, 10 µg $ml^{-1}$ meropenem, or 8 µg $ml^{-1}$ colistin. The reference measurement for the biofilm model was carried out with a 2-stack agarose disk (Assembly B), while the reference measurement for the mucoid biofilm sample was from a 3-stack agarose disk (Assembly D).

It is important to note that the extra agarose disc added to the assembly of the mucoid biofilms is not responsible for the significantly reduced diffusion noticed in gentamicin and colistin. This was proven by using an assembly model consisting of 3 stacks of agarose disks, with comparable heights to that of the test samples as a reference measurement for this assay (b).

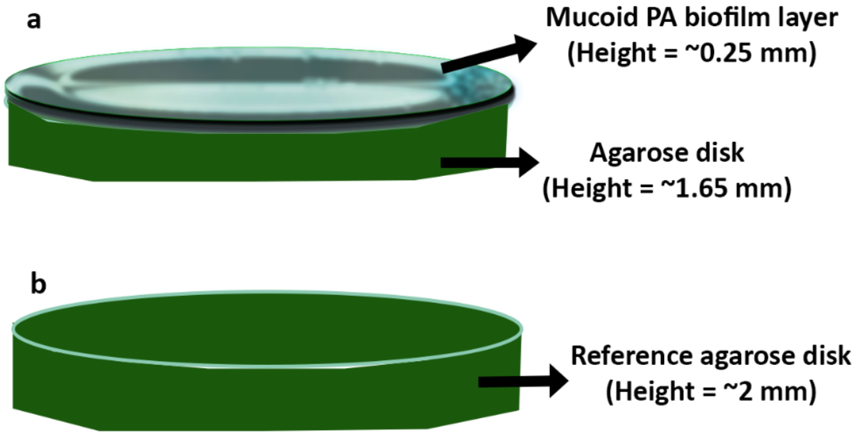


**Figure S6**. Schematic representation of a) mucoid PA biofilm growth on an agarose disk and b) reference disk, respectively. To create a biofilm layer, a mucoid PA culture ($OD_{600}$ = 0.1) was pipetted onto agarose disks in Petri dishes and incubated for 1 h to allow bacterial attachment. Excess bacteria were removed by washing three times with 1 ml TSB. Subsequently, 50 µl of TSB was added to 145 mm Petri dishes containing agarose disks, ensuring the surface remained exposed to air while receiving nutrients from below. Bacteria-loaded agarose disks were incubated for 3 days at 37 °C for biofilm development on the surface of the disks.

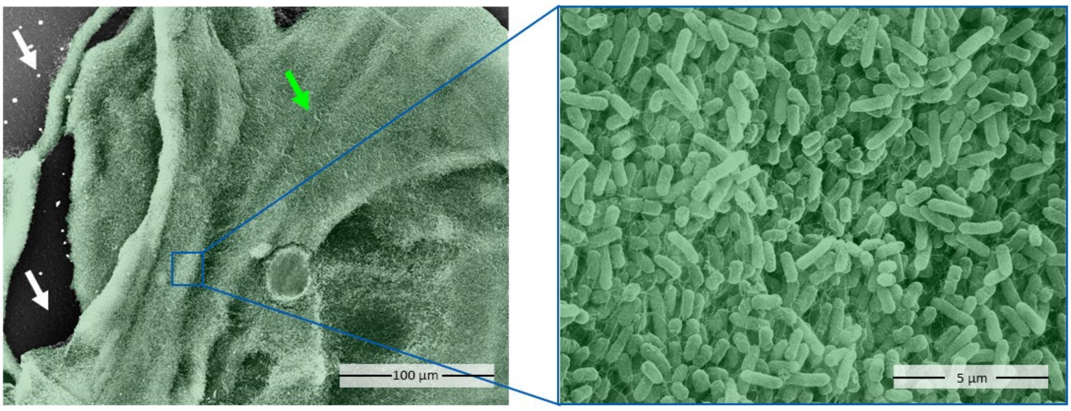


**Figure S7**. SEM Image of mucoid PA biofilm layer on agarose disk. White arrow – agarose. Green arrow - biofilm layer. The biofilm layer is manually coloured green.

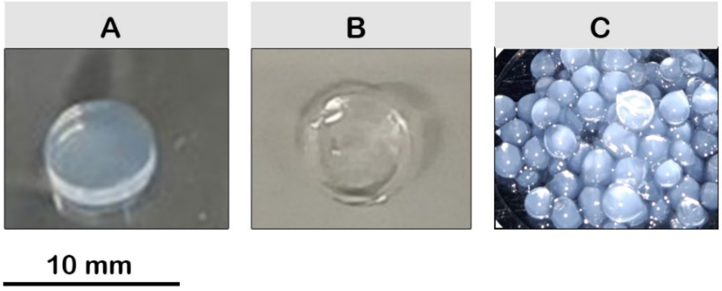


**Figure S8.** Alginate and agarose models. **a)** acetylated alginate **b)** non-acetylated alginate, **c)** agarose, **d)** acetylated alginate beads
